# Starburst amacrine cells amplify optogenetic visual restoration through gap junctions

**DOI:** 10.1101/2020.08.11.246686

**Authors:** Yusaku Katada, Hiromitsu Kunimi, Naho Serizawa, Deokho Lee, Kenta Kobayashi, Kazuno Negishi, Hideyuki Okano, Kenji F. Tanaka, Kazuo Tsubota, Toshihide Kurihara

## Abstract

Ectopic induction of optogenetic actuators, such as channelrhodopsin, is a promising approach to restoring vision in the degenerating retina. However, the cell type-specific response of ectopic photoreception has not been well understood. It is limited to obtain efficient gene expression in a specifically targeted cell population by a transgenic approach. In the present study, we established a murine model with high efficiency of gene induction to retinal ganglion cells (RGCs)- and amacrine cells using an improved tetracycline transactivator-operator bipartite system (KENGE-tet system). To investigate the cell type-specific visual restorative effect, we expressed the channelrhodopsin gene into RGCs and amacrine cells using the KENGE-tet system. As a result, enhancement in the visual restorative effect was observed to RGCs and starburst amacrine cells. In conclusion, a photoresponse from amacrine cells may enhance the maintained response of RGCs and further increase/improve the visual restorative effect.

## Introduction

Inherited retinal degeneration is one of the major causes of vision loss. More than 2 million people worldwide are blind due to this disease ^1^, and there is still almost no effective treatment. Previous studies have reported visual restoration effects in animal models by various molecules such as optogenetic actuators ^2–9^. In addition, clinical trials have also started using channelrhodopsin 2 (RST-001, ClinicalTrials.gov Identifier: NCT01648452) and Chrimson R (GS-030, ClinicalTrials.gov Identifier: NCT03326336), with gene transduction into retinal ganglion cells (RGCs) by intravitreal injection of recombinant adeno-associated virus (rAAV). Although the visual reconstruction effects by optogenetic gene transfer (such as channelrhodopsin-2 into RGCs^2^ and ON-bipolar cells ^3,5,6,8,10^) have been shown, interactions between the other types of cells in the retinal neural circuits in optogenetic visual restoration have not been well understood yet. Channelrhodopsin 2 conductance was reported to be 50–250 fS ^11^, indicating the need for sufficient gene expression to control the membrane potential. In this study, we employed a tetracycline-controllable gene expression system (tet system) ^12^ in which the amount of gene expression has been much improved (KENGE-tet system) ^13^. Furthermore, we established sufficient gene induction in RGCs and amacrine cells. To investigate cell-type-specific visual restorative effects, channelrhodopsin2(E123T/T159C) was ectopically expressed into those cells by KENGE-tet system. As a result, we revealed that amacrine cells may play an essential role in retinal neuronal circuits via gap junctions to enhance the optogenetic visual restorative effect.

## Results

### Gene induction of RGCs and SACs in M4-YC and RGCs in 5B-YC

We employed two different mouse lines that express the gene encoding the tetracycline transactivation (tTA) protein under the control of a cell-type-specific promoter, muscarinic acetylcholine receptor 4 (*Chrm4*)^14^ or 5-hydroxytryptamine (5-HT, serotonin) 5B receptor (*Htr5b*)^13^ control region: *Chrm4*-tTA or *Htr5b*-tTA. These mice were further crossed with another transgenic mouse line containing the yellow Cameleon-Nano 50 (YC) fluorescent gene connected into the downstream of the tet operator (tetO) promoter ^15^. The YC gene expression was induced only by the presence of tTA protein in the double transgenic mice (*Chrm4*-tTA::tetO-YC (M4-YC) or *Htr5b*-tTA::tetO-YC (5B-YC)) (Figure S1A, B). The expression of YC was observed in the double transgenic mouse retina with a fluorescence microscope. In the M4-YC mouse retina, we identified the expression of YC in RGCs (Figure 1A-D, Figure S2A-E) and amacrine cells (Figure 1A-D, Figure S1E-F) with *in vivo* fluorescence microscopy and in sections. In the 5B-YC mouse retina, we identified the expression of YC in RGCs (Figure 1E-H) and the corneal stromal layer (Figure S2G-M) with *in vivo* fluorescence microscopy and in sections. In both lines, no expression was found in displaced amacrine cells, only RGCs were labeled in the ganglion cell layer, and the percentage of YC-positive cells in RGC marker RBPMS ^16^-positive cells was 35.4% (n=5) in M4-YC and 36.5% (n=5) in 5B-YC from RGCs, which were equivalent (Figure 1I-L, O-R, Figure S2M-Q). All of the amacrine cell expression in the M4 line was choline acetyltransferase (ChAT) staining-positive and consisted of starburst amacrine cells (SACs), and 31.1±2.1 % (n=3) of ChAT-positive cells were YC-positive (Figure 1M-T, Figure S3 A-L).

**Figure 1.**
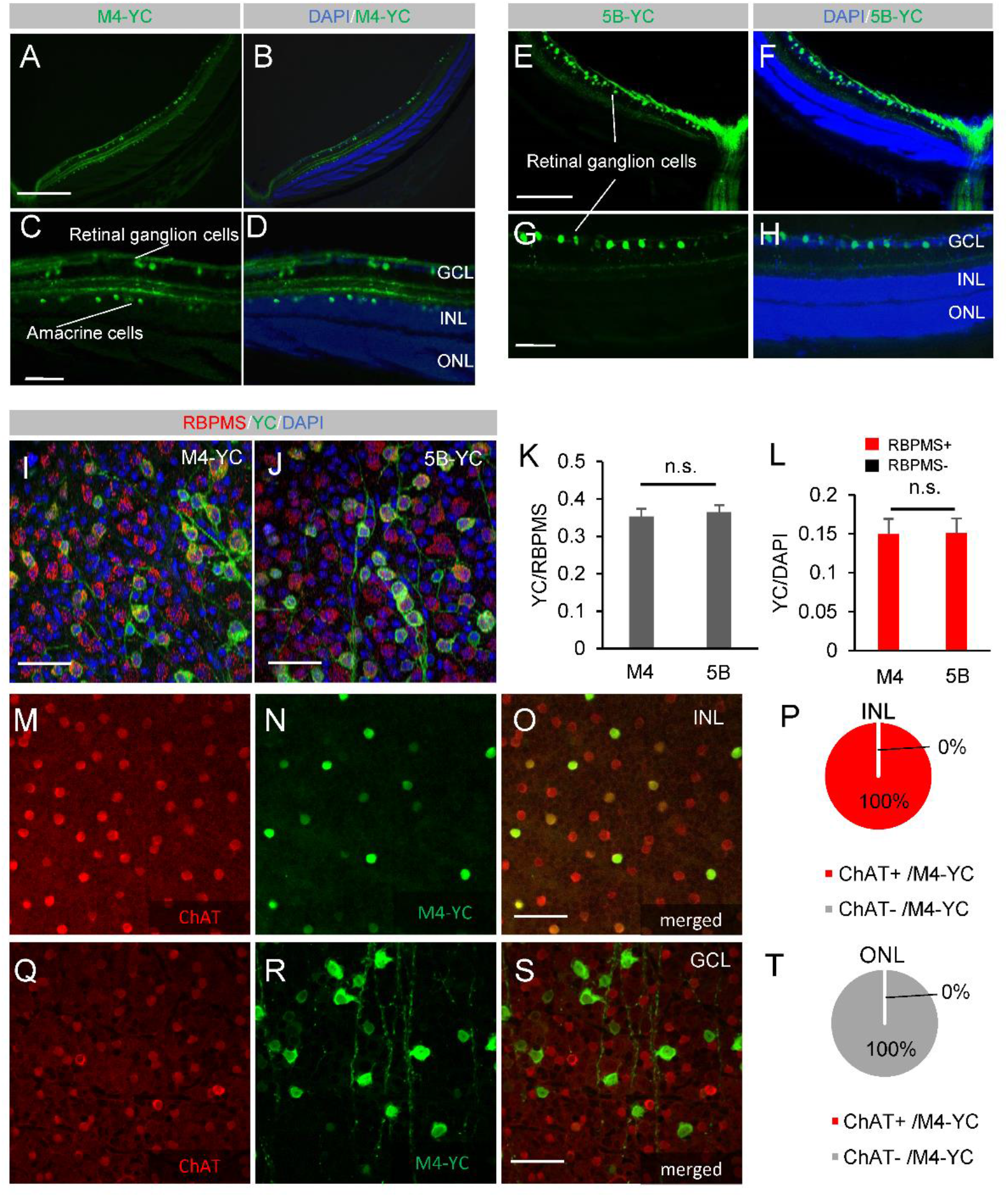
Gene induction of RGCs and SACs in M4-YC and RGCs in 5B-YC. In the M4-YC mouse retina, we identified the expression of YC (green) in RGCs and amacrine cells within sections (A-D). In the 5B-YC mouse retina, we identified the expression of YC (green) in RGCs in sections (E-H). Coexpression of the RGC marker RBPMS in flat-mounted retinas of M4-YC(I) and 5B-YC(J). Percentage of YC-positive cells in RBPMS-positive (K) or DAPI-positive cells (L) and RBPMS-positive cells in YC-positive cells (L) in both lines from confocal flat-mounted GCL (n = 3 retinas each). Regions were chosen in each quadrant, and we obtained RBPMS, DAPI-positive, YC-positive, and co-labeled cells. Coexpression of the starburst amacrine cell marker ChAT in flat-mounted retinas of M4-YC in INL (M-O) and GCL (Q-S). Percentage of ChAT-positive cells in YC-positive cells in M4-YC mice from INL (P) and GCL (T) (n = 3 retinas each). Error bars represent the SEMs. INL, inner nuclear layer; GCL, ganglion cell layer; ONL, outer nuclear layer. Scale bar: 50 µm in (E). 100 μm in (B-D), (H), (I), (K), (M), (Q), and (R). 400 µm in (N). 500 µm in (F), (G), (O), and (P).

As a visual restoration model, the *Chrm4*-tTA and *Htr5b*-tTA line were crossed with tetO-channelrhodopsin2(E123T/T159C) (tetO-ChR2) mice ^17^ (Figure S1C). *Chrm4*-tTA::tetO-ChR2 (M4-ChR2) drove ChR2 expression in both RGCs and amacrine cells and *Htr5b*-tTA::tetO-ChR2 (5B-ChR2) only in RGCs in the retina. In the retinas of these double transgenic mice, photoreceptor degeneration was induced by intraperitoneal injection of N-methyl-N-nitrosourea (MNU) ^18^. MNU is an alkylating agent that causes DNA methylation in the O6 position of guanine, resulting in apoptosis, and it has been widely used to induce a pharmacological animal model of retinitis pigmentosa (RP). Two weeks after 75 mg/kg MNU administration into the mice at the age of 8 weeks, the outer nuclear layer containing photoreceptors was largely absent (Figure S4A, B), and the light-evoked response from photoreceptors was not detected by electroretinography (ERG) (Figure S4C).

TUNEL assay was performed to determine whether there was any neurotoxicity-induced cell death in the retina due to Channel rhodopsin expression, however, we could not observe any cell death in the retina (Figure S4D).

### M4-ChR2 mouse shows higher visual restorative effect

To evaluate the function of the channelrhodopsin ectopically induced in the transgenic mouse retinas, we performed multi-electrode array (MEA) recording, which can record extracellular potentials of RGCs (Figure S4E). As a result of photoreceptor degeneration induced with MNU treatment, the control retina without channel-rhodopsin expression (tetO-ChR2) showed no response from RGCs detected (Figure 2A). In contrast, the M4-ChR2 (Figure 2B) and 5B-ChR2 (Figure 2C) retinas showed obvious light-induced responses. After filtering with the Gaussian function (Figure 2D, E), the maintained response after the peak was significantly greater in M4-ChR2 mice than in 5B-ChR2 mice (Figure 2F), which indicates that the light-evoked ON response in RGCs could be modified through the response from amacrine cells.

**Figure 2.**
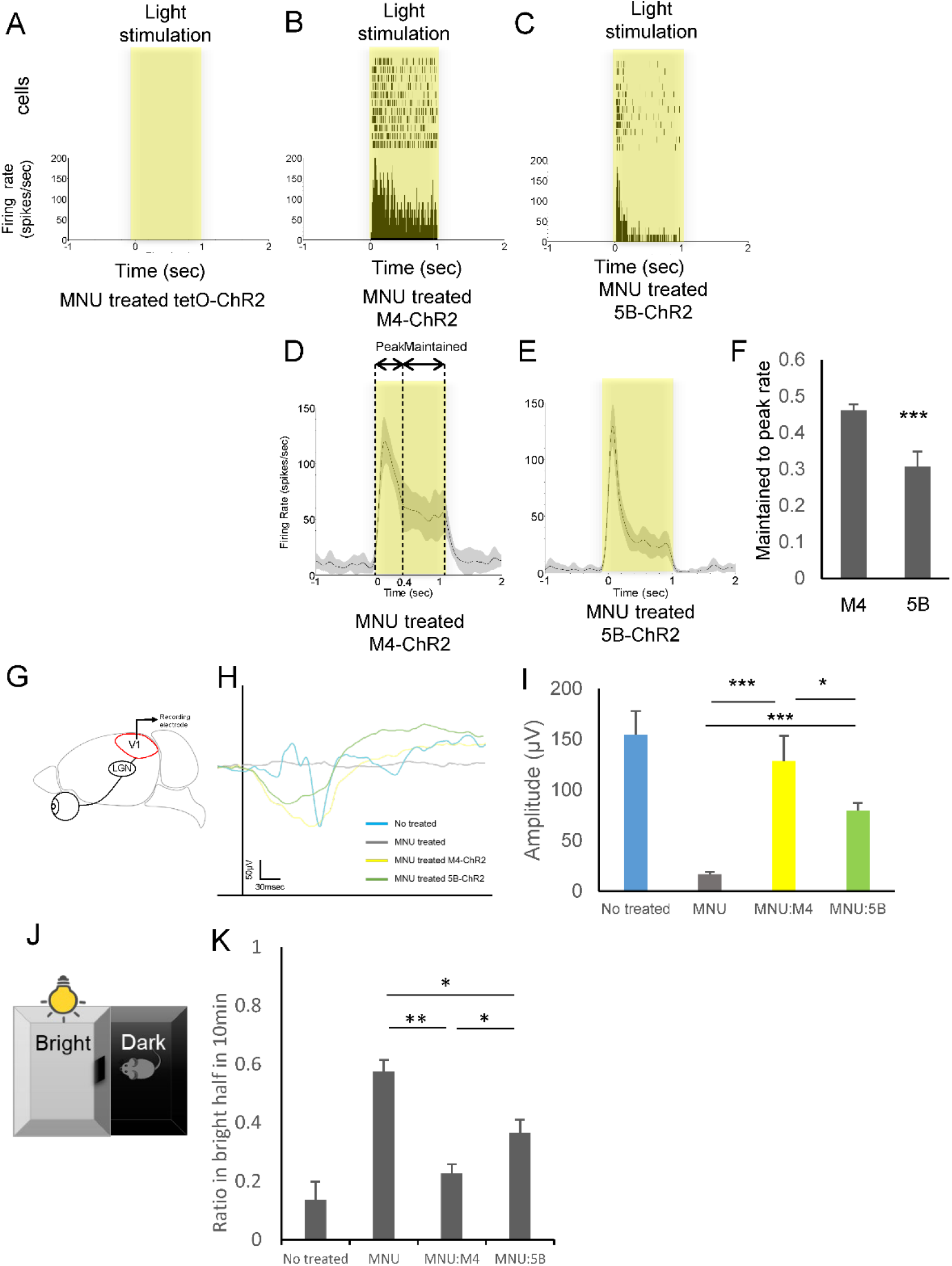
M4-ChR2 mouse shows a higher visual restoration effect. (A-C) Raster plots and peri-stimulus time histogram (PSTH) of light stimulation from single RGCs of MNU-treated tetO-ChR, M4-ChR2, and 5B-ChR2 mice. Light intensity was 13.6 log photons/cm^2^/s, and the duration was 1.0 seconds. (D, E) The averaged rate histogram after filtering with the Gaussian function from M4-ChR2 and 5B-ChR2 mice. At least 10 trials were conducted for each cell. The gray areas around the averaged traces represent the SEM. (F) Maintained-to-peak ratio of the spiking responses recorded. The maintained time frame is 0.4 to 1.0 seconds from light stimulation as shown in H (n = 7 retinas, 164 cells in MNU-treated M4-ChR2 mice, and n = 4 retinas, 127 cells in MNU-treated 5B-ChR2 mice). Error bars represent SEMs. ***p < 0.001. Student’s 2-tailed t-test. (G) Schematic image of VEP measurement. (H) Representative VEP traces from MNU-injected and control mice. (I) The average amplitude of the VEPs in the control tetO-ChR mice (n = 4), MNU-treated tetO-ChR mice (n = 8), M4-ChR2 mice (n = 14), and 5B-ChR2 mice (n = 12) at 10 weeks of age. It was stimulated with a light stimulus intensity of 100-ms pulses of white LED 4,000 cds/m2. Signals were low-pass filtered at 300 Hz and averaged over the 60 trials. (J) Schematic image of the LDT. Mice were tested in a 30 × 45 × 30-cm box containing equally sized bright (200 lx at ground level) and dark chambers connected by a 5 × 5-cm opening, across which the mice could move freely. Visible mice feel uneasy in bright places, so staying time in the bright half gets shorter. (K) % time in bright half at 10 min of control (tetO-ChR2 mice) (n = 4), MNU-injected tetO-ChR2 mice (n = 8), MNU injected M4-ChR2 mice (n = 15), and MNU-injected 5B-ChR2 mice (n = 8) measured from LDT. All error bars represent the SEMs. *p < 0.05, **p < 0.01, ***p < 0.001. One-way ANOVA testing.

To investigate whether light reception on the retina is transmitted to the visual cortex, we then examined visual evoked potentials (VEPs) from the visual cortex (Figure 2G). The output from RGCs is sent through the axons of RGCs (the optic nerve) to the lateral geniculate nucleus (LGN) of the thalamus, which is a region of the mesencephalon, from the LGN to the primary visual cortex (V1) in the occipital lobe of the cerebral cortex. VEPs were not detected in the control tetO-ChR2 mice with MNU treatment (Figure 2H, I). In contrast, VEPs were observed in both M4-ChR2 and 5B-ChR2 mice treated with MNU (Figure 2H, I). In response to the light stimulus at 4000 cds/m^2^, the average of the VEP amplitude in MNU-treated M4-ChR2 mice was significantly higher (143.6 μV; n=12, P=0.03) than in MNU-treated 5B-ChR2 (79.6 mV; n = 12) and the same level as in the mice without MNU treatment (155 μV; n=4). However, the shapes of the waveforms were irregular in both models compared to controls, and their physiological roles are unknown. Thus, there is a limitation in this direct comparison.

To validate the model system, we also examined the MEA, ERG and VEPs of M4-ChR2 and 5B-ChR2 in the absence of retinal degeneration (no MNU treatment and no mutation). As a result of MEAs, the peak response was significantly reduced in both lines and further reduced with MNU treatment (Figure S4F). A similar trend was observed for the maintained response, whereas it was maintained in the M4 line (Figure S4G). This finding suggests that channelrhodopsin gene induction into healthy RGCs interferes with the physiological retinal light response. In addition, in the control mice, not only the ON response but also the OFF response was confirmed. In contrast, the OFF response was significantly lost in the M4 line (Figure S4I). It is known that SACs in the INL are connected to off bipolar cells ^19^, which might be related to this change. As a result of ERG and VEPs, both lines tended to have shorter latencies, especially in photopic examinations (Figure S5E, I, L, O), likely because of the channelrhodopsin response to strong light ^9^ and its photoreception in RGCs resulting in short latency ^2^. Although the amplitude of both ERG and VEP tended to be small in the 5B line (Figure S5C, G, M, N), these measurements were not significant, and there was no significant change in the shape of the waveforms.

Next, light-dark transition testing (LDT) was performed to investigate whether ectopic expression of channelrhodopsin in the degenerative retinas may lead to behavioral changes according to visual restoration. Rodents tend to stay in dark places according to their visual function since they are nocturnal and feel uneasy in bright environments (Figure 2J), while the visually disturbing MNU-treatment (Figure 2K) resulted in almost half of the staying time in bright and dark places (ratio in bright half at 10 min was closer to 0.5). In contrast, both the M4-ChR2 and 5B-ChR2 mouse lines with each MNU-treatment showed decreased staying time in bright places, indicating that visual restoration was confirmed in these models. Furthermore, M4-ChR2 mice showed significantly higher visual restorative effects than 5B-ChR2 mice (Figure 2K).

We also examined the optokinetic response (OKR) to investigate whether light receptivity in the retina restored by ectopic channelrhodopsin expression could lead to central reflex movement output. In OKR, a rotating striped pattern was displayed to the head-fixed mice to induce eye movements, and the velocity was measured to evaluate the integrity of the subcortical reflex circuitry of the mice (Figure S6A). OKR was not detected in the mice with MNU administration except for the M4 line (4.80 deg/sec, n=6) (Figure S6B).

This outcome seems to be due to lack of ChR2 expression in SACs, which play a key role in OKR ^20^, in the retinas of 5B-ChR2 mice. We also examined *rd1* mice as a genetic animal model of RP. These mice have a nonsense mutation in the *Pde6b* gene leading to rapid degeneration of rods, photoreceptors followed by loss of cones ^21,22^. We used blind *rd1* mutation mice at the age of 10-12 weeks for the following experiments. In the cases of *rd1* mutant mice, no detectable OKR was observed in any combinations with each transgenic line, including M4-ChR2. To investigate the dissociation of OKR between the MNU-treatment and *rd1* models, the retinal thickness was compared under the same conditions above. As a result, the total retinal thickness of *rd1* mice (78.3 μm; n=9) was significantly thinner than that of MNU-treated mice (97.2 μm; n=10) evaluated on optical coherence tomography (OCT) (Figure S6C, D). This outcome was thought to be because the inner retinal layer was thinner of the *rd1* mouse (Figure S6E). Indeed, the inner nuclear layer of the *rd1, rd1*;5B-ChR2 and *rd1*;M4-ChR2 mice was significantly thinner than that of the MNU-treated mice (Figure S6F) and YC-positive cells in the inner retinal layer of *rd1*;M4-ChR2 tended to be fewer than those in the inner retinal layer of MNU-treated mice (Figure S6G), suggesting that SACs in the inner layer may be degenerated. This result suggests that residual inner retinal thickness and its durability might be potential limitations of optogenetic gene therapy in inherited retinal disease patients clinically.

### Acetylcholine and gap junctions are involved in the maintained response

The M4-ChR2 mouse line, which showed the transgene expression in amacrine cells in addition to RGCs, showed a higher maintained response and more effective visual restoration (Figure 2) than the 5B-ChR2 mice. Therefore, we investigated the neurocircuit pathway responsible for the enhanced response due to amacrine cells using neurotransmitter blockers on the MEA recording. Administration L-2-amino-4-phosphonobutyric acid (L-AP4), which is an agonist for group III metabotropic glutamate receptors, including mGluR6 working as a blocker of retinal ON-bipolar cells, did not show significant changes in either MNU-treated M4-ChR2 (Figure 3A-C) or 5B-ChR2 (Figure 3D-F) retinas, which indicates that the photoresponse was not derived from photoreceptors. In contrast, administration of mefenamic acid (MFA) or carbenoxolone, an inhibitor of gap junctions, induced a significant decrease in the maintained response but not the peak of the response in the MNU-treated M4-ChR2 retina (Figure 3G-I, Figure S7A-C). This change was recovered after washout and not observed in the 5B-ChR2 retina (Figure S7D-F). SACs release GABA and acetylcholine ^23^. Thus, we also examined inhibitors of GABA receptors, bicuculline for GABA-A, CGP 52432 for GABA-B and (1,2,5,6-tetrahydropyridin-4-yl) methylphosphinic acid (TPMPA) for GABA-C receptors, and cholinergic antagonists, mecamylamine for nicotinic and atropine for muscarinic acetylcholine receptors. As a result, no significant change was observed with the administration of GABA receptor blockers (Figure S7G-O) and mecamylamine (Figure S7P-R), but atropine administration showed a similar decrease in maintained response and peak response in the MNU-treated M4-ChR2 mice (Figure 3J-L). In addition, to determine whether this change may occur via bipolar cells, an AMPA receptor antagonist, 6-cyano-7-nitro-quinoxaline-2,3-dione (CNQX), and an NMDA receptor antagonist, D(-)-2-amino-5-phosphonopentanoic acid (D-AP5), were administered. As a result, no statistically significant difference was observed, although there was a tendency toward a decrease in the maintained response (Figure S7S-U). SACs form synapses with bipolar cells, RGCs, and other amacrine cells ^23–25^. It is also known that amacrine cells are directly connected with RGCs through gap junctions ^26,27^, regulating neural circuits in the retina ^28^. Therefore, these results indicated that SACs enhanced the maintained response of RGCs directly or indirectly through muscarinic acetylcholine and gap junctions.

**Figure 3.**
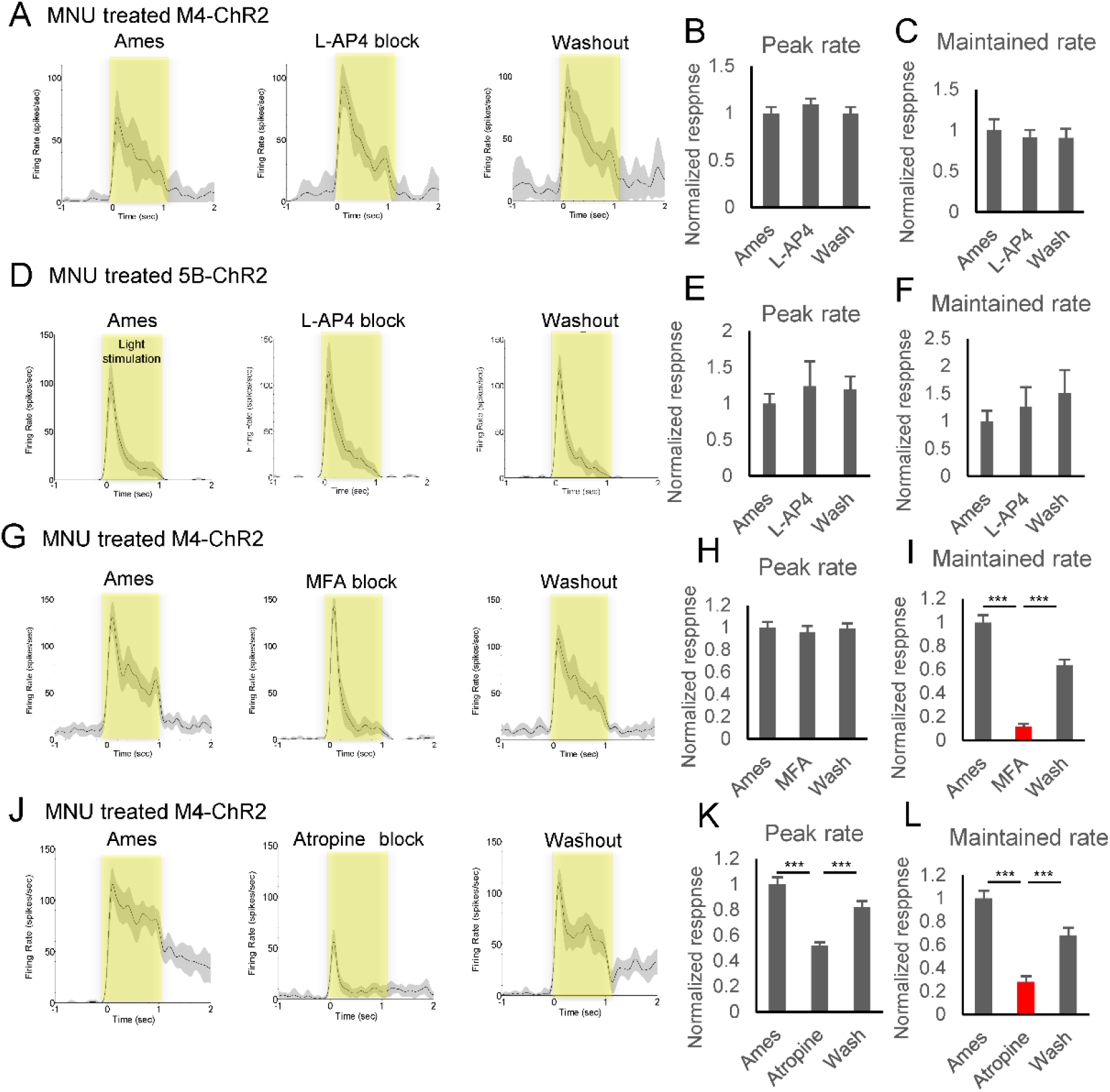
Acetylcholine and gap junctions were involved in the maintained response. (A, D, G, J) Mean ± SEM of exemplar cell response firing rate recorded during normal Ames’ medium superfusion (left), in synaptic block (middle), and after washout (right). MNU-treated M4-ChR2 mice with L-AP4 block (n = 3 retinas, 47 cells) (A), MNU-treated 5B-ChR2 mice with L-AP4 block (n = 3 retinas, 45 cells) (D), MNU-treated M4-ChR2 mice with MFA block (n = 3 retinas, 85 cells) (G), MNU-treated M4-ChR2 mice with atropine block (n = 3 retinas, 33 cells) (J). The gray areas around the averaged traces represent the SEM. (B, C, E, F, H, I, K, L) Averaged normalized peak firing rate and maintained rate. The maintained time frame is 0.4 to 1.0 seconds from light stimulation. Light intensity was 13.6 log photons/cm^2^/s. All error bars represent the SEM. ***p < 0.001. One-way ANOVA and Tukey’s test.

MEA showed a visual restoration effect induced by ectopic ChR2 expression in the degenerative retina (Figure 2). These photoresponses obtained from MEA were all ON responses, as previously reported ^2,29,30,31.^ Several slow photoresponses obtained from RGCs were considered to have an ipRGC origin ^32^, and they were excluded from the data. The MNU-treated M4-ChR2 retina showed a significant more enormous response in the maintained time phase compared to 5B-ChR2. As a result of a neurotransmitter blocking test (Figure 4), inhibition of muscarinic acetylcholine and gap junctions decreased the sustained response of only M4-ChR2 retinas. The difference between M4-ChR2 and 5B-ChR2 was the presence or absence of ChR2 gene expression in SACs. Therefore, it was suggested that photoresponse from SACs enhanced the maintained response of RGCs, enhancing the visual restoration effect via muscarinic acetylcholine and gap junctions.

**Figure 4.**
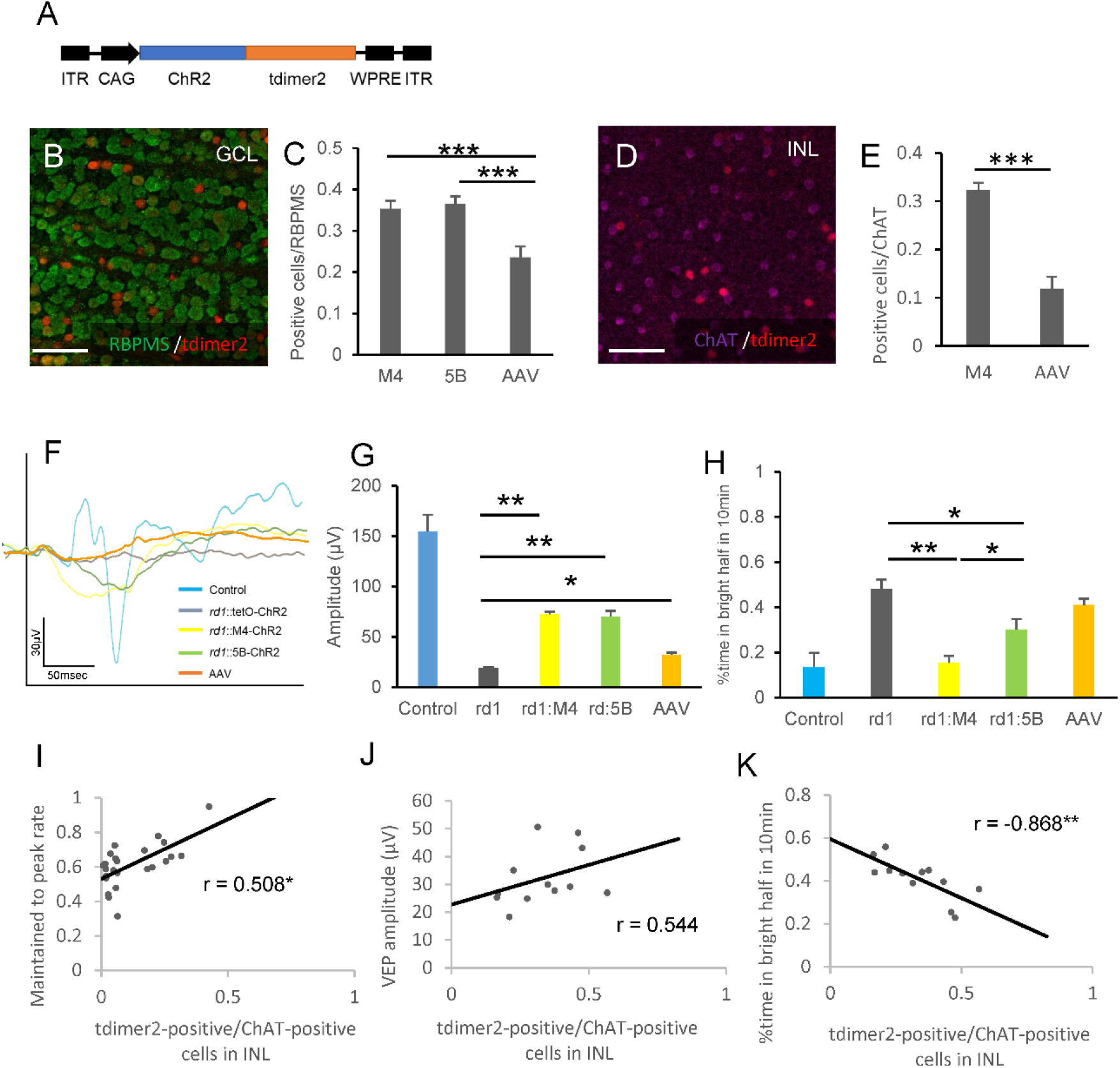
Induction efficiency into SACs tended to affect visual restoration. (A) The rAAV2–CAG-ChR2–tdimer2–WPRE expression cassette: ITR, inverted terminal repeat; WPRE, woodchuck hepatitis virus posttranscriptional regulatory element. (B-E) Coexpression of the RGC marker RBPMS (B) and starburst amacrine cell marker ChAT (D) and tdimer2 in flat mounted retinas of rAAV2–CAG–tdimer2– WPRE treated retinas in *rd1* mice. (C, E) Percentage of YC-positive cells in M4-YC and 5B-YC mice and tdimer2-positive cells in rAAV-treated mice in RBPMS-positive (C) or ChAT-positive cells (E) from confocal flat mounted retina (n = 3 retinas each). Regions were chosen in each quadrant, and we obtained RBPMS, ChAT-positive, YC/tdimer2-positive, and co-labeled cells. (F, G) Average VEP traces (F) and quantification of its amplitudes (G) from control (tetO-ChR2) (n = 3), *rd1*::tetO-ChR2 (n = 6), *rd1*;M4-ChR2 (n = 12), *rd1*;5B-ChR2 (n = 12) and rAAV treated *rd1*::tetO-ChR2 mice (n = 12) at 10 weeks of age. It was stimulated with 100-ms pulses of white LED 4,000 cds/m^2^ light stimulus intensity. Signals were low-pass filtered at 300 Hz and averaged over the 60 trials. (H) % time in bright half at 10 min in control (tetO-ChR2) (n = 3), *rd1*::tetO-ChR2 (n = 6), *rd1*;M4-ChR2 (n = 12), *rd1*;5B-ChR2 (n = 12) and rAAV-treated *rd1*::tetO-ChR2 mice (n = 12) measured from LDT. (I-K) Correlation between transfection efficiency into SACs (tdimer2-positive cells/ChAT-positive cells in INL) and maintained to peak rate (I) (n = 24), VEP amplitude (J) (n = 24) and % time in bright half in LDT (K) (n = 12). All error bars represent the SEMs. INL, inner nuclear layer; GCL, ganglion cell layer. Scale bars, 50 µm in (B), (D), n.s.: not significant, *p < 0.05, **p < 0.01, ***p < 0.001. unpaired t-test (E),Games-Howell test (C, G, H), Pearson’s correlation coefficient (I-K).

### Induction efficiency into SACs tends to affect visual restoration

Finally, to investigate its clinical applicability, the effect of SACs on visual restoration was examined using rAAV2 which was approved for gene therapy ^33^. We injected rAAV2-CAG-ChR2-tdimer2 intravitreally into *rd1* mice and performed MEA, VEPs, and LDT (Figure 4A, Figure S8A, B). In addition, we counted the number of transfected RGCs and SACs and analyzed their relevance to visual restorative effects. As a result of immunohistochemical labeling using RBPMS, the infection efficiency to RGCs was 24%, which was equivalent to previous reports (21-23%) ^2,34,35^, and significantly less than those of our transgenic mice (Figure 4B, C). The efficiency of SACs was 12%, which was also less than that of transgenic mice (Figure 4D, E). As a result of MEA recording, the peak response in the rAAV model was equivalent to that of both transgenic lines (Figure S8C). The maintained response and its rate were equal to that of the M4 line (Figure S8D, E), perhaps because the input from various cell types other than SACs is also included in the rAAV model. The increased maintained response may be affected by gene transfer to cells other than SACs, but similar characteristics to the M4 line have been obtained. Each average of the VEP amplitude in the *rd1*;M4-ChR2, *rd1*;5B-ChR2, and rAAV-treated *rd1* mice in response to the light stimulus at 4000 cds/m^2^ was 72.7 μV (n=10), 70.4 μV (n=12) and 32.2 μV (n=12), respectively. These responses were significantly higher than those in *rd1*;tetO-ChR2 mice (19.1 mV; n=8) but smaller than those in the 8-week-old wild-type C57BL/6J mice (155 μV; n=4) (Figure 4F, G). There was no significant difference between *rd1*;5B-ChR2 and *rd1*; M4-ChR2 mice, and the VEPs of rAAV treated *rd1* mice were smaller than those of them (Figure 4F, G). Even in the LDT results, the restorative effect of rAAV-treated *rd1* mice tended to be lower than that of transgenic mice (Figure 4H), perhaps due to its infection efficiency or the higher maintained response might not lead directly and simply to better visual restoration. In addition, the correlation between the infection efficiency of SACs in each rAAV-treated *rd1* mouse and the visual restorative effect was investigated. The results showed a significant, positive correlation between the number of transfected SACs in the inner nuclear layer and the maintained to peak ratio in MEA (Figure 4I), a positive correlation between the VEP amplitudes (Figure 4J), and a significant, negative correlation between the time spent in the bright half in LDT (Figure 4K). This outcome suggests that the maintained response in MEA is derived from SACs and that, clinically, the inclusion of SACs, rather than only RGCs, which is the primary target of optogenetic therapy, is effective for visual restoration. Although there is a limitation because infection efficiency is a confounding factor, there was no correlation between the number of transfected RGCs in the ganglion cell layer and the maintained to peak ratio in MEA (Figure S8F). The correlation of the number of transfected SACs in the inner nuclear layer with the VEP amplitudes (Figure 4J) and the time spent in the bright half in LDT (Figure 4K) was stronger than that with infection efficiency into RGCs (Figure S8G, H).

## Discussion

In this study, we established transgenic mouse lines inducing gene induction in specific retinal cells using the tet-system (Figure S1). We identified that gene induction occurred in RGCs and SACs under the control of a muscarinic acetylcholine receptor *Chrm4* promoter. In contrast, serotonin (5-HT) *Htr5b* receptor control region led to induction only in RGCs in the retina (Figure 2). Among acetylcholine receptors expressed by all types of neurons in the retina ^36–41^, the *Chrm4* muscarinic receptor is expressed in RGCs and amacrine cells ^36,42^. All of the amacrine cell expressions in the M4 line consisted of type-a SACs (Figure 1M-T, Figure S3 A-L). There are two subtypessubtypes of SACs, type-a, and type-b (displaced SACs)^43,44^. This would be the first report of a transgenic line specific to only type-a but not type-b SACs. Approximately 30% of the type-b SACs were expressed in the M4 line, and no noticeable morphological differences were observed between positive and negative SACs. The 5-HT *Htr5b* receptor is expressed in rodents but not in humans ^45^. Although several types of 5-HT receptors are expressed in the mouse retina ^46^, the retinal distribution of the *Htr5b* receptor has not been previously described to our knowledge. We found that these gene inductions in both mouse retinas are useful for examining the functions of RGCs and SACs.

Accumulating data have shown that the visual restoration strategy induced by optogenetic genes is a promising therapy for degenerative retinal diseases. Most amacrine cells are inhibitory neurons in the vertebrate retina, which have not been regarded much in the elementary visual restoration target. This study showed that SACs increased the maintained response through gap junctions and contributed to enhancing the visual restorative effect. In particular, since the restorative effect of LDT and OKR was more significant than that of VEPs, it might contribute to sustained behavior and direction recognition rather than transient response, consistent with the role of the maintained response in RGCs ^47^. Including the results using rAAV, in the current situation, viral delivery in primates is limited ^29^, and gene transfer involving amacrine cells, for example, using ubiquitous promoters, might be more effective in visual restoration than limiting RGCs in the clinical setting.

In conclusion, this study suggests that the optogenetic photoresponse from SACs enhances the maintained response, which further enhances the visual restoring effect. These results may provide the basis for advanced visual restoration in the future.

## Methods

### Key Resources Table

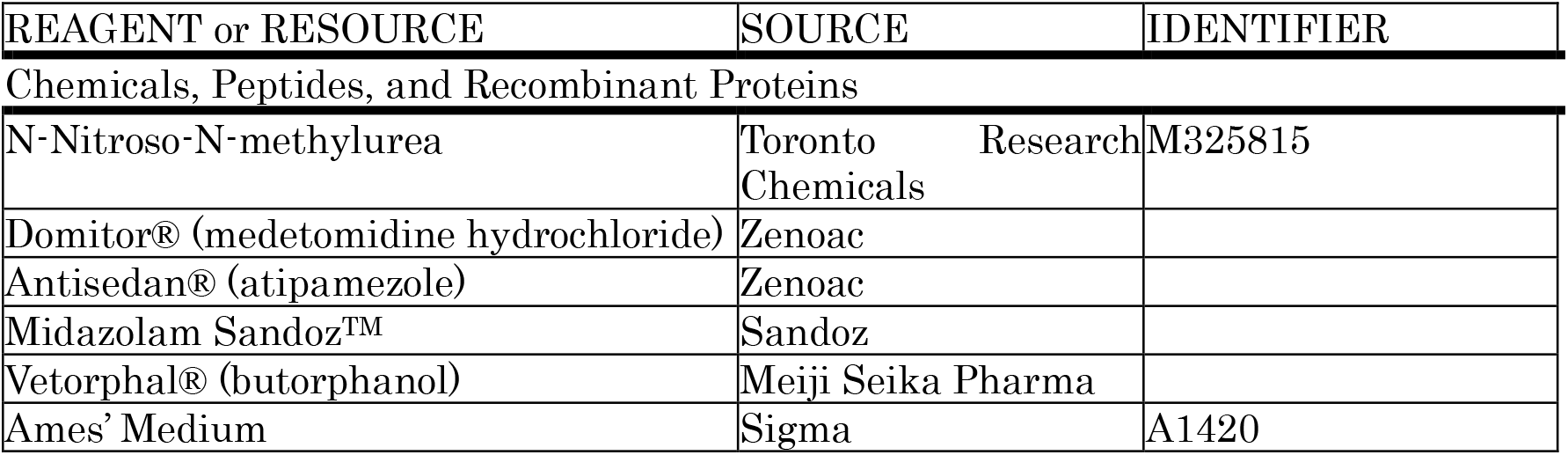
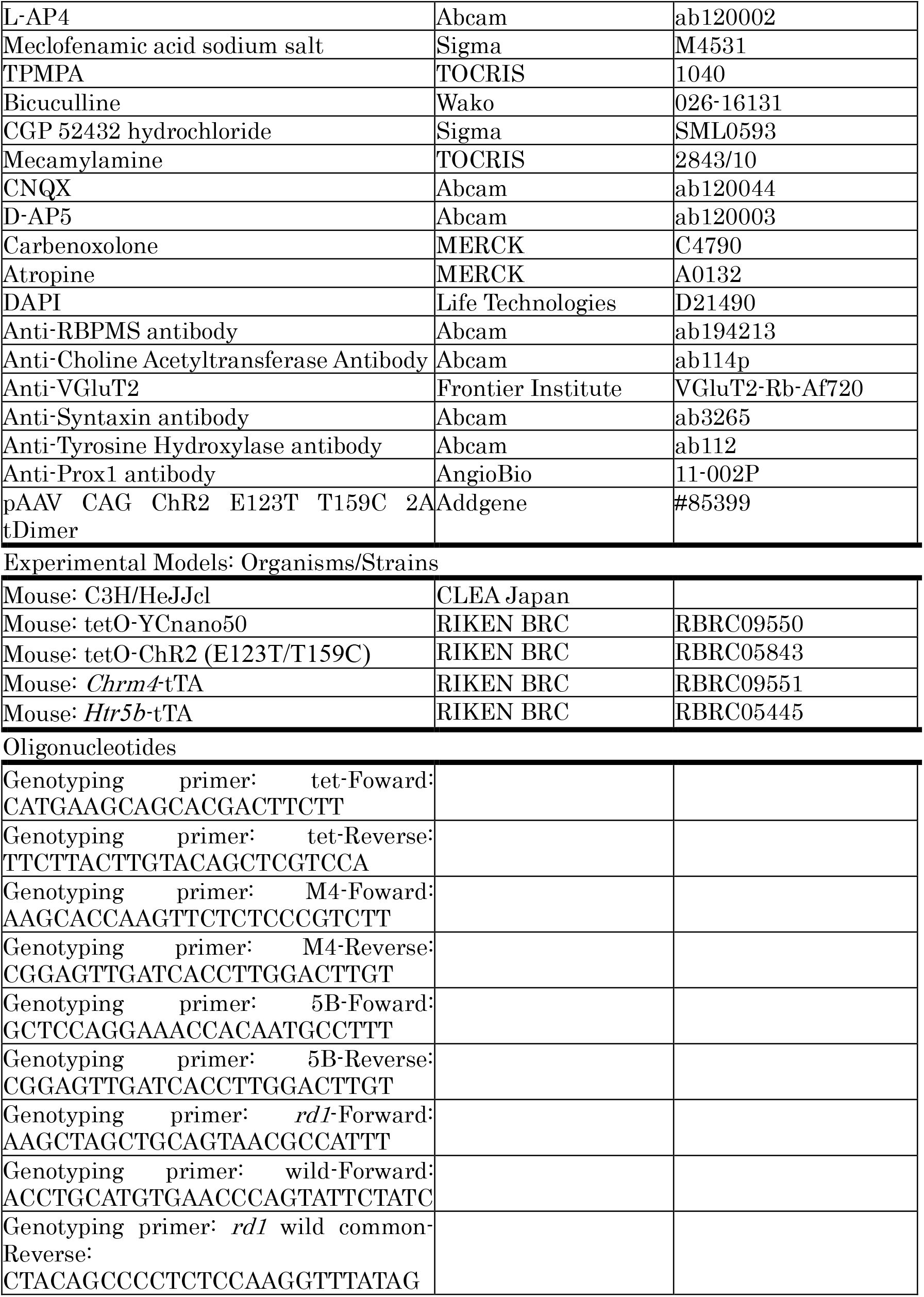

### Animals

Transgenic mice used for the experiments and their genotyping protocols were generated as previously reported^13^. Mice homozygous for the retinal degeneration alleles Pde6b^*rdl*^ (C3H/HeJJcl, *rd1*) and WT C57BL/6J were obtained from CLEA Japan, Inc. Animals were maintained under 12-h light:12-h-dark conditions. For animals bred in-house, littermates of the same sex were randomized to experimental groups. Mice used for the experiments were heterozygous for the tTA and tetO genes and homozygous for the *rd1* gene. All of the animal experiments were conducted under protocols approved by the Institutional Animal Care and Use Committee of Keio University School of Medicine.

tetO-YC (RBRC09550), tetO-ChR2 (RBRC05843), *Chrm4*-tTA (RBRC09551), and *Htr5b*-tTA (RBRC05445) are available from RIKEN Bioresource Center in Japan.

Study approval: All of the animal experiments were conducted under protocols approved by the Institutional Animal Care and Use Committee of Keio University School of Medicine (#2808).

### Immunohistochemistry

The protocol for immunohistochemistry as previously described^48^. The retinas were incubated in phosphate-buffered saline (PBS) with 1% Triton X-100 and 0.5% Tween 20 for 1 h at room temperature and in 4% BSA for 1 h at room temperature and then incubated overnight at 4 °C with primary antibodies: RBPMS (1:500, Abcam, Cambridge, UK), ChAT (1:100, Abcam), VGLUT2 (1:100, Frontier Institute, Hokkaido, Japan), syntaxin (1:100, Abcam), tyrosine hydroxylase (1:100, Abcam) and Prox1 (1:100, AngioBio, San Diego, CA, USA) in blocking buffer. Secondary anti-rabbit, mouse, and goat IgG, conjugated with Alexa TM488, TM594, and 633, respectively (1:1000; Molecular Probes), were applied for 1 h at room temperature. The retinas then were flat-mounted, and the sections were mounted on slide glass. The TUNEL assay was performed based on our previous reports^49,50^.

### Vector production and purification

pAAV-CAG–ChR2(E123T/T159C)-tdimer2–WPRE (Addgene, Watertown, MA) was used for the current study. Type 2 serotypes of rAAV vectors were prepared using the AAV Helper Free Packaging System (Cell Biolabs, San Diego, CA, USA). The serotypes were produced in HEK293 cells using a helper virus-free system and were purified using two CsCl2 density gradients and titrated by quantitative polymerase chain reaction. Final preparations were dialyzed against PBS and stored at −80°C.

### Virus injection

The mice were anesthetized with a combination of midazolam, medetomidine, and butorphanol tartrate at doses of 4 mg/kg, 0.75 mg/kg, and 5 mg/kg of body weight and placed on a heating pad that maintained their body temperatures at 35–36°C throughout the experiments. An aperture was made subsequent to the limbus through the sclera with a 30-gauge disposable needle, and a 33-gauge unbeveled blunt-tip needle on a Hamilton syringe was introduced through the scleral opening into the vitreous space for intravitreal injections and introduced through the scleral opening along the scleral interior wall into the subretinal space for subretinal injections. Each eye received 1 µl of vehicle (PBS) or vector at a titer of 2.0 × 10^11^ vg/ml.

### Multielectrode array recordings (MEA)

All of the MEA procedures were performed under dim red light. The mice were anesthetized and euthanized by quick cervical dislocation. Following enucleation, the retina was dissected at room temperature in Ames’ medium bubbled with 95% O2/5% CO2 (A 1420; Sigma-Aldrich). The separated retina was placed on a cellulose membrane, and RGCs were directed to the electrode and was gently contacted against MEA (MEA2100-Systems; Multi-Channel Systems, Reutlingen, Germany) under suction pressure. During the experiment, the retinas were continuously perfused with Ames’ medium bubbling at 34 ° C. at a rate of 1-2 ml/min. Recorded signals were collected, amplified, and digitized using MC Rack software (Multi-Channel Systems). Retinas were perfused for 30 min in darkness before recording responses. A 470-nm blue LED light was used at 1.0 × 10^16^ `photons·cm−2·s−1 stimulus. L-(+)-2-amino-4-phosphonobutyric acid (L-AP4,` ab120002; Abcam), (1,2,5,6-tetrahydropyridin-4-yl) methylphosphinic acid (TPMPA, Cat. No. 1040; Tocris Bio-Science), meclofenamic acid sodium salt (MFA, M4531; Sigma-Aldrich), atropine (A0132; MERCK), carbenoxolone (C4790; MERCK), Bicuculline (026-16131; Wako), CGP52432 (SML0593; Sigma) and Mecamylamine (2843/10, TOCRIS) were newly diluted to 20 µM, 20 µM, 100 µM, 20 µM, 100 µM, 10μM, 20 µM and 20 µM respectively. Stimulation was presented for 1 second at 60-second intervals. Signals were filtered between 200 Hz (low cutoff) and 20 kHz (high cutoff). A threshold of 40 µV was used to detect action potentials, and action potentials from individual neurons were determined via a standard expectation–maximization algorithm using Off-line Sorter software (Plexon, Dallas, TX, USA). The results were plotted using NeuroExplorer software (Nex Technologies Colorado Springs, CO, USA). Maintained-to-peak amplitude ratio was calculated by dividing the maintained response amplitude in maintaining time frame (0.4 to 1.0 seconds after light stimulation) by the peak amplitude (this ratio quantifies the sustenance of the response).

### ERG analyses

Scotopic ERG was recorded according to our previous report ^48^. Animals were dark-adapted for 12 h and prepared under dim red illumination. The pupils were dilated with a mixed solution of 0.5% tropicamide and 0.5% phenylephrine (Mydrin-P; Santen, Osaka, Japan). Then, the mice were anesthetized with a combination of midazolam, medetomidine, and butorphanol tartrate at doses of 4 mg/kg, 0.75 mg/kg, and 5 mg/kg of body weight, respectively, and were placed on a heating pad that maintained their body temperature at 35–36°C throughout the experiments. The ground electrode was a subcutaneous needle in the tail, and the reference electrode was placed subcutaneously between the eyes. The active contact lens electrodes (Mayo, Inazawa, Japan) were placed on the corneas. Recordings were performed with a PuREC acquisition system (Mayo). Responses were filtered through a bandpass filter ranging from 0.3 to 500 Hz to yield a-and b-waves. White LED light stimulations of 10.0 log cd-s/m^2^ were delivered via a Hemisphere LS-100 Stimulator (Mayo).

### VEP analyses

The measuring electrodes for VEP analyses were placed more than one week before the measurement. The mice were anesthetized with a combination of midazolam, medetomidine, and butorphanol tartrate at doses of 4 mg/kg, 0.75 mg/kg, and 5 mg/kg of body weight, respectively. The animals were placed in a stereotaxic holder. A stainless-steel screw (M1.0×6.0 mm) inserted through the skull into both visual cortex (1.5 mm laterally to the midline, 1.5 mm anterior to the lambda), penetrating the cortex to approximately 1 mm, served as a measuring electrode.

At the measurement time, the mice were anesthetized again with the same doses. Visual stimuli were generated by a white LED placed 3 cm from the eye. It was stimulated with 100-ms pulses of white LED 4000 cds/m^2^ light stimulus intensity. Signals were acquired and analyzed with a PuREC acquisition system (Mayo). Signals were low-pass filtered at 300 Hz and averaged over the 60 trials.

### OKR recording system

Protocols for eye movement recording and visual stimulation were previously described ^51,52^. Eye movements were recorded from both eyes of each animal separately. During the recording, the contralateral eye was covered by aluminum foil. The head of the mouse was fixed to an experimental steel board by the head-mounted stick for the LIM lens frames. The reflected images through a hot mirror (43957-J, Edmund) were recorded using an infrared CCD camera (BS-GV200, Libraly Inc., Tokyo, Japan). The images of the eye movements were processed and analyzed using software (Move-tr/2D, Libraly Inc., Tokyo, Japan). The sampling rate of the image was 200 Hz. The center of the pupil was detected by the software. We calculated the speed of the eye movements on two-dimensional images and converted them to angular speeds using the AL of each eye. The spatial frequency was set as 0.125 cycles/degree, and the temporal frequency of the visual stimulus was 1.5 Hz. The motion onset delay (MOD) was set as 333 msec. Continuing the MOD, sinusoidal grating started to move clockwise in 5 sec. The intervals of the visual stimulus were 60 sec. Eye movements were recorded three times for each experiment to exclude shaking images due to excessive body movements. Average velocities of the eye movements were calculated in the slow speed phase of their nystagmus.

### LDT recording

Mice were tested in a 30 × 45 × 30-cm box containing equally sized light and dark chambers connected by a 5 × 5-cm opening via which mice could move freely. The bright half of the box was illuminated from above by a white fluorescent light with an intensity of 200 lux measured at the floor level. The animals were placed in the bright half, and movement was recorded (HD Pro Webcam C920, Logitech, Lausanne, Switzerland). A trial lasted 10 min, and then the testing apparatus was dismantled and cleaned with 70% ethanol. Videos were analyzed using ANY-maze tracking software and were validated by comparison with manual analysis. Time spent in the bright half was recorded.

### Preparation of whole-mount samples and cryosections of retinas

Enucleated eyes were fixed for 20 min in 4% paraformaldehyde (PFA) in PBS and then dissected as previously described ^53^. The obtained tissues were post-fixed overnight in 4% PFA and stored in methanol at –20°C. Cryosections of retinas (12 mm) were prepared as previously described ^54^ after the eyeballs were immersed overnight in 4% PFA. Retinal sections were observed using a confocal microscope (LSM710; Carl Zeiss, Jena, Germany).

### OCT Imaging

The thickness of the retina was analyzed by an SD-OCT system (Envisu R4310; Leica, Wetzlar, Germany) tuned for mice. The imaging protocol entailed a 3 mm×3 mm perimeter square scan sequence producing a single en-face image of the retina through a 50-degrees field of view from the mouse lens, following mydriasis. The en-face image consisted of 100 B-scan tomograms, with each B-scan consisting of 1000 A-scans. The retinal thickness of 150 µm from the optic disc of each quadrant was measured.

### Quantification and Statistical Analysis

All of the results are expressed as the mean ± SEM. The averaged variables were compared using the Student’s 2-tailed t-test and the Kruskal Wallis one-way ANOVA test. P-values of less than 0.05 were considered statistically significant.

## Data and Software Availability

Raw MEA spike data were sorted offline to identify single units using Offline Sorter software (version 4.4.0)(Plexon). Spike-sorted data were analyzed with NeuroExplorer 5 software (version 5.115) (Nex Technologies).

## Author Contribution

Y.K. and T.K. designed the research, wrote the manuscript, and performed the retinal histology, MEA, ERG and VEP recordings, and LDT experiments. K.K. performed AAV production. Y.K. and H.K. performed OKR recordings. N.S. performed MEA. D.L. performed a TUNEL study. Y.K. performed data processing and analysis. K. T, K.N., H.O, and T.K. made critical revisions to the manuscript. T.K. supervised the research.

## Acknowledgments

We would like to thank Prof. Amane Koizumi for critical comments to the manuscript. Y.K. is supported by grants from the Keio University Doctorate Student Grant-in-Aid Program.

T.K. is supported by Grants-in-Aid from Takeda Science Foundation and the Keio University Medical Science Fund.

