## Supplemental materials for "Starburst amacrine cells amplify optogenetic visual restoration through gap junctions"

### Supplemental Information for

#### Starburst amacrine cells amplify optogenetic visual restoration through gap junctions in the murine retina

### Supplemental Figures

#### Figure S1. KENGE-tet system and tetO-Yellow cameleon

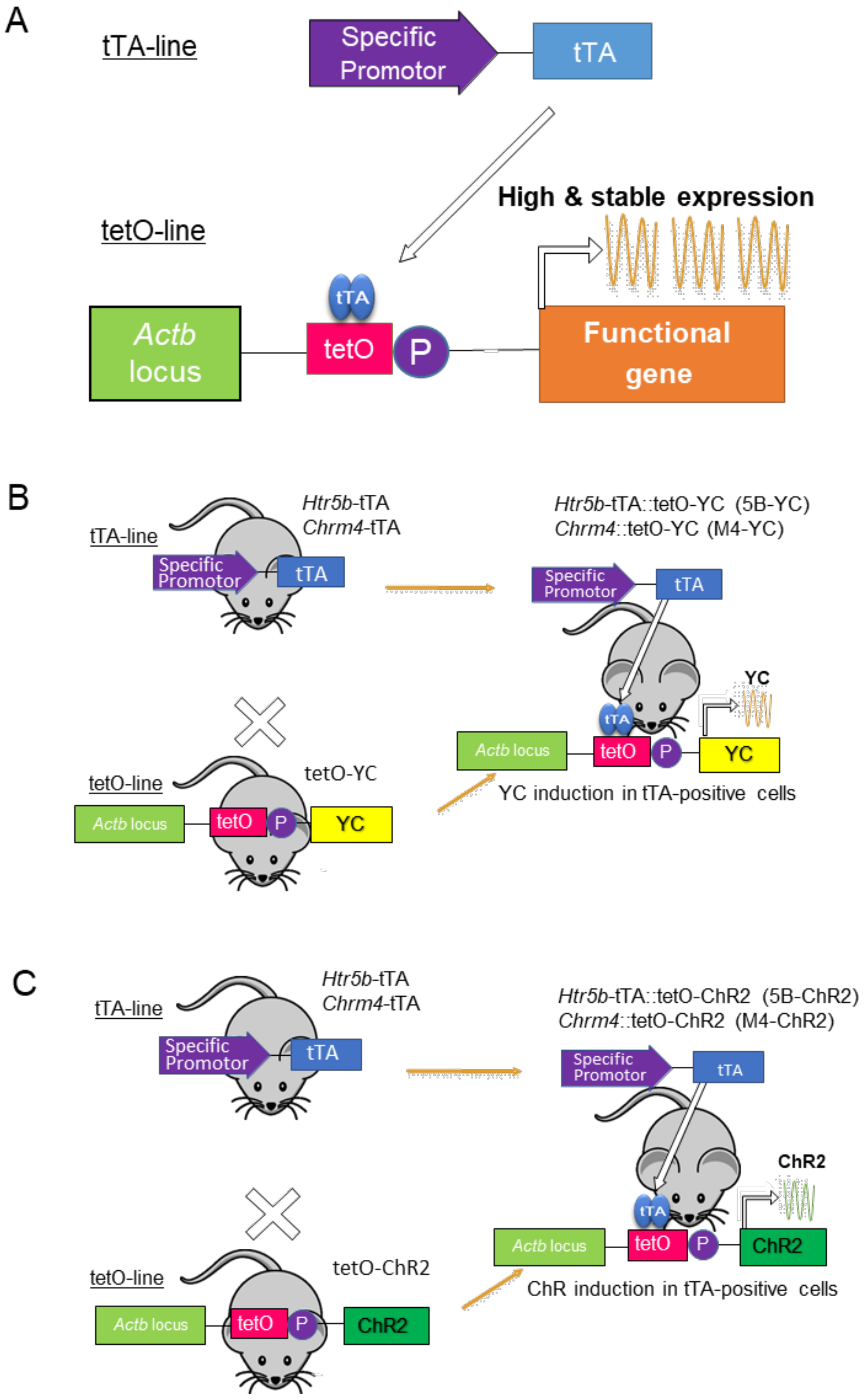

(A) KENGE-tet system. tTA is expressed under the control of cell type-specific promoters. tTA transactivates the tetO promoter, and the functional protein is induced in a cell-type-specific manner. (B, C) Two different mouse lines are employed that express the gene encoding tTA protein under the control of a cell-type-specific promoter, muscarinic acetylcholine receptor M4 or serotonin receptor 5B control region. These mice were further crossed with another transgenic mouse line containing a YC fluorescent gene connected into the downstream of the tetO promoter. The YC gene expression was induced only by the presence of tTA protein in the double transgenic mice (M4-YC or 5B-YC)(B). Next, as a visual restoration model, the tTA line was crossed with tetO-ChR2. (C) tTA drove ChR2 expression in RGC.

**Figure S2. Expression of Yellow cameleon in M4-YC and 5B-YC**

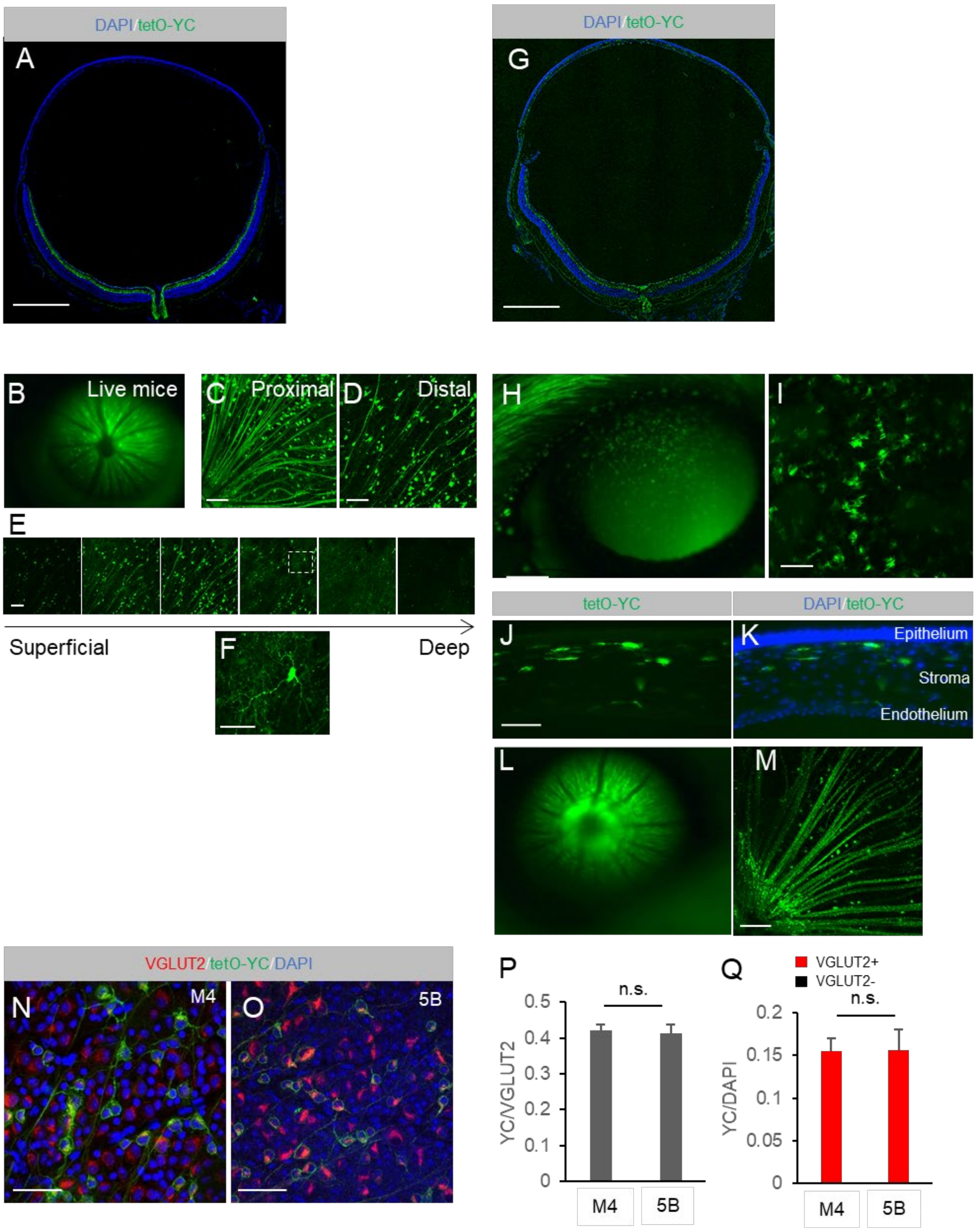

In the M4-YC mouse retina, we identified the expression of YC (green) in RGC and amacrine cells within sections (A), with in vivo fluorescence microscopy (B), and with flat mounted retina (C-F). In the 5B-YC mouse retina, we identified the expression of YC in RGC and the corneal stromal layer with in sections (G, J, K), in vivo fluorescence microscopy (H, I, L), and the flat mounted retina (M). Coexpression of the RGC marker VGLUT2 in flat mounted retina of M4-YC (N) and 5B-YC (O). Percentage of YC-positive cells in VGLUT2-positive (P) or DAPI-positive cells (Q) and VGLUT2-positive cells in YC-positive cells (Q) in both lines from confocal flat mounted GCL (n = 3 retinas each). Regions were chosen in each quadrant, and we obtained VGLUT2, DAPI-positive, YC-positive, and co-labeled cells. Error bars represent the SEM. Scale bar: 50 μm in (F), (N) and (O). 100 μm in (C-E), (I), (J) and (M). 1,000 μm in (A) and (G).

**Figure S3. Immunostaining of amacrine cells in M4-YC mice**

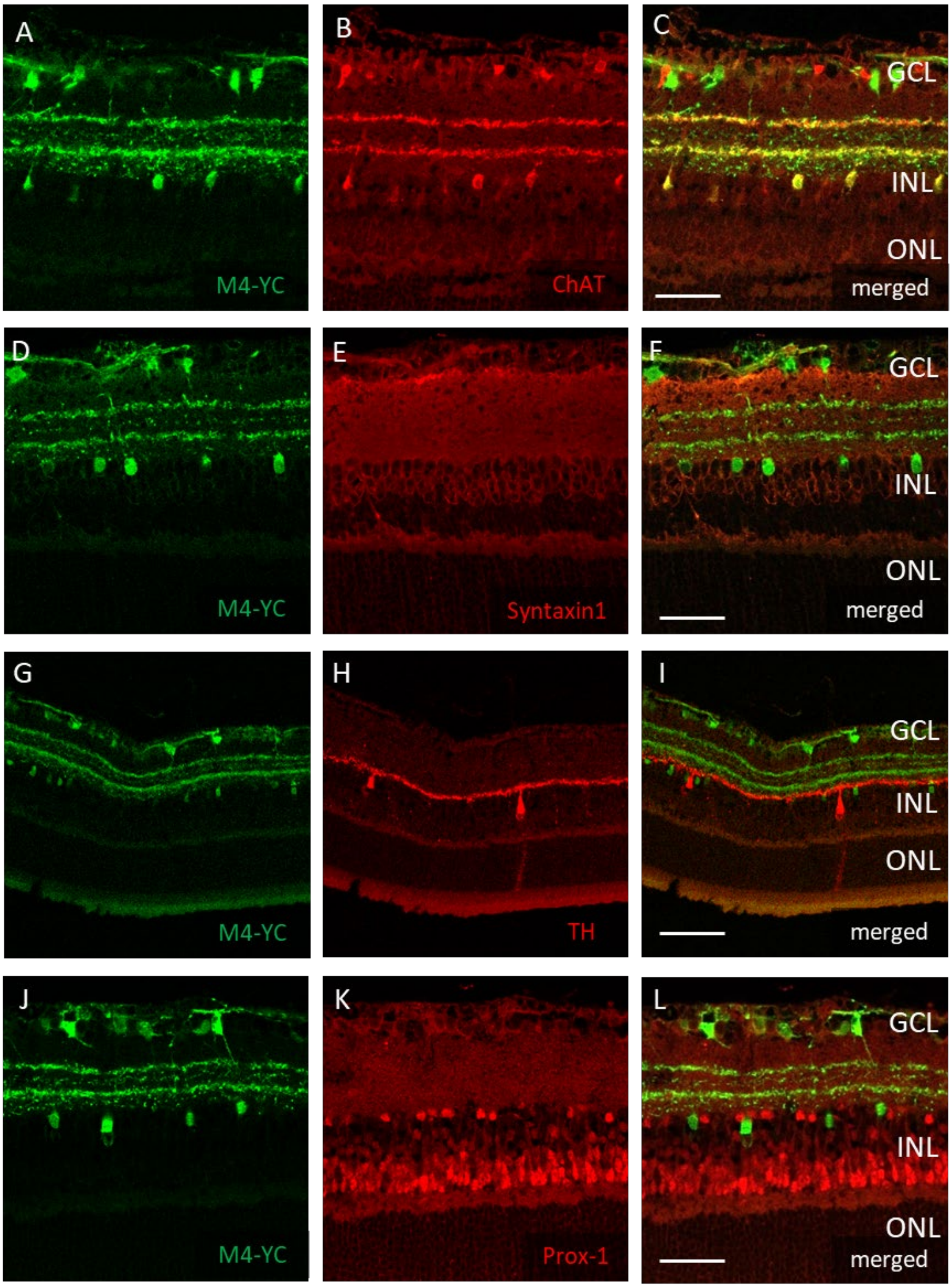


Immunohistochemistry on the transverse retinal cryosection of M4-YC mouse labeled with choline acetyltransferase (ChAT), a starburst amacrine marker (A-C), syntaxin1, a pan-amacrine cell marker (D-F), tyrosine hydroxylase (TH), a dopaminergic amacrine marker (G-H) and Prox-1, an AII amacrine marker (J-L). GCL, ganglion cell layer; INL, inner nuclear layer; ONL, outer nuclear layer. Scale bars, 50 µm in A-L.

**Figure S4. The maintained response was retained regardless of photoreceptor degeneration in M4-ChR2M4-ChR2 mice.**

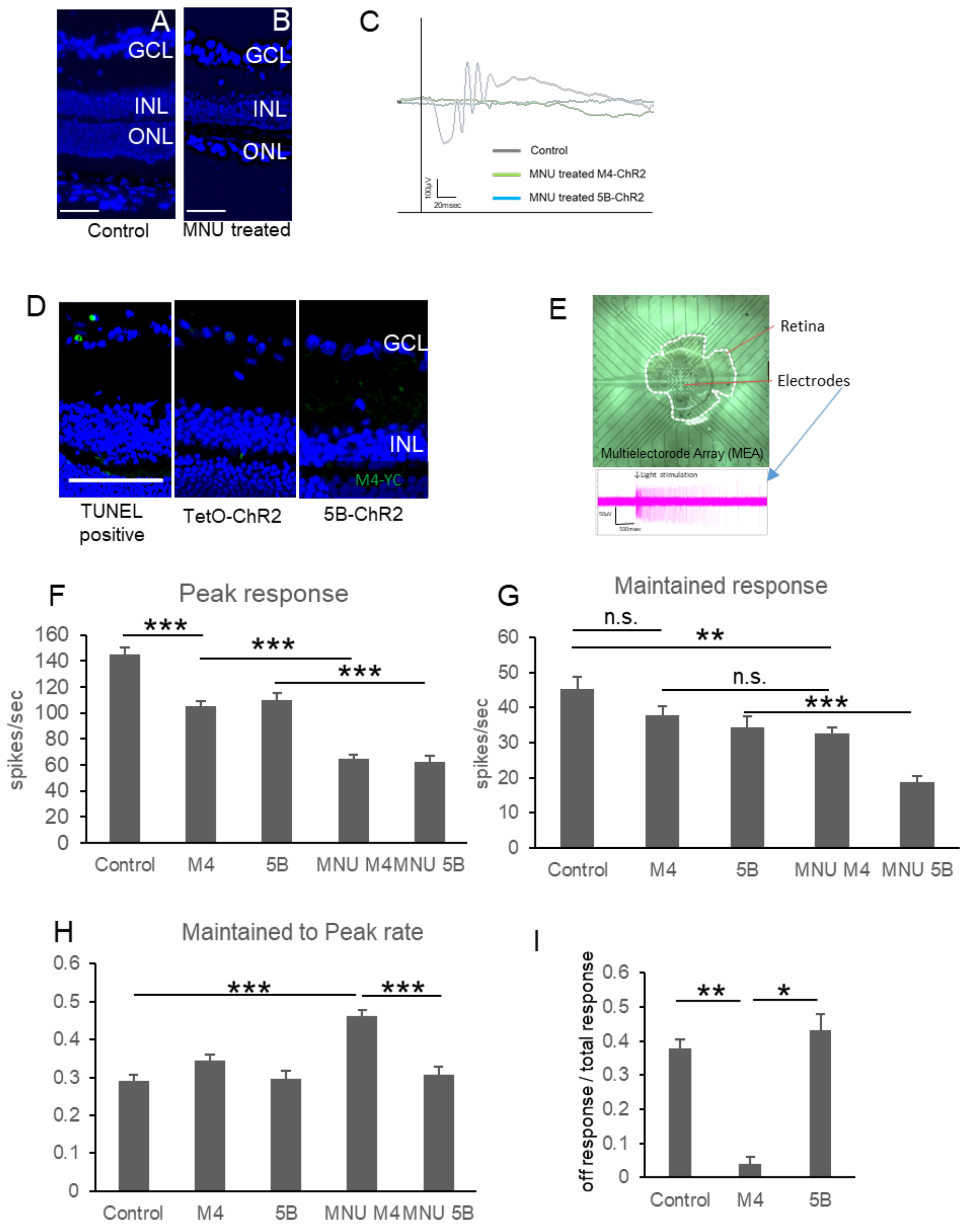


(A, B) Comparison of retinal sections after intraperitoneal administration of MNU. MNU was used to produce a photoreceptor degeneration model. Two weeks after the MNU injection, DAPI nuclear counter-stain is shown in blue. Scale bar: 50 µm. (C) Representative

ERG traces from MNU-injected and control mice. White LED light stimulation of 10.0 log cd-s/m$^2$ was delivered. (D) Sections of a representative TUNEL assay from a positive control (A murine model of retinal Ischemia/Reperfusion injury; left), a negative control mouse (tetO-ChR2; middle), and ChR2 expression sample (5B-ChR2; right). DAPI nuclear counter-stain is shown in blue and TUNEL staining is shown in green. Scale bar: 100 µm. (E) Image of MEA. It can measure the extracellular potential of RCGs in contact with the electrode ex vivo. (F-G) Comparison of peak response (F), maintained response (G) and maintained to peak rate (H) from MEA recordings among control (tetO-ChR2; n = 3 retinas, 112 cells), M4-ChR2 (n = 3 retinas, 62 cells), 5B-ChR2 (n = 3 retinas, 48 cells), MNU-treated M4-ChR2M4-ChR2 (n = 3 retinas, 164 cells) and MNU-treated 5B-ChR2 (n = 3 retinas, 117 cells) mice. All ON responses were used for analysis in mice without retinal degeneration. (I) Comparison of ratio of off response to total light response from MEA recording. Error bars represent SEMs. *$p < 0.05$, **$p < 0.01$, ***$p < 0.001$. Games-Howell test.

**Figure S5. Ectopic channelrhodopsin induction had no significant effect on ERG or VEP.**

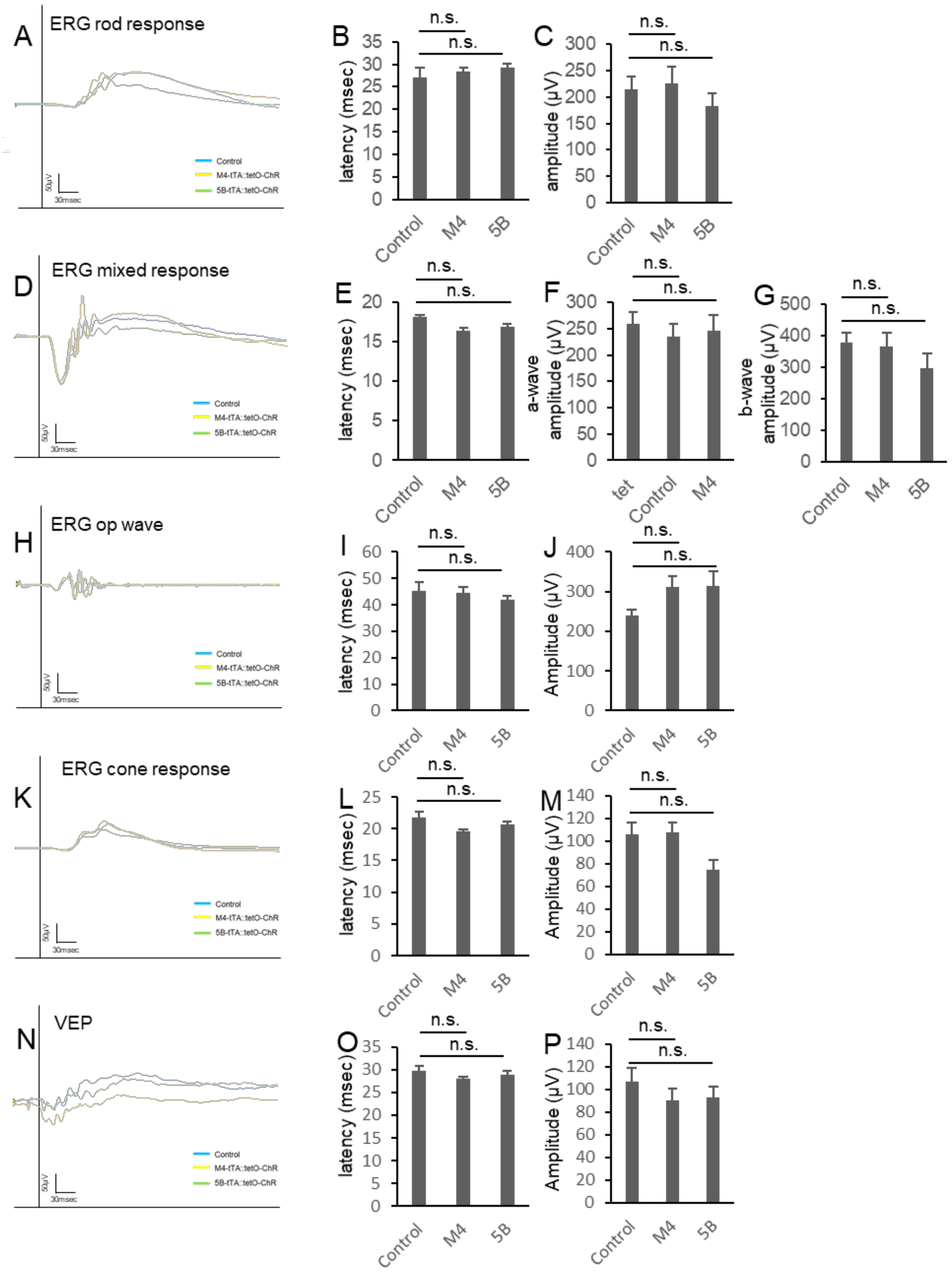

Representative ERG waveforms from ERG rod response (A), mixed response (D), oscillatory potentials (H), cone response (K) and VEPs (N). Quantification of latency (B, E, I, L, O) and amplitude (C, F, G, J, M, P) in each protocol in control (tetO-ChR2; n = 6), M4-ChR2M4-ChR2 (n = 8), and 5B-ChR2 (n = 8) mice. Error bars represent SEMs. n.s., not significant. Games-Howell test.

**Figure S6. Thinning of INL in the genetic model of retinal degeneration attenuating the restoration effect.**

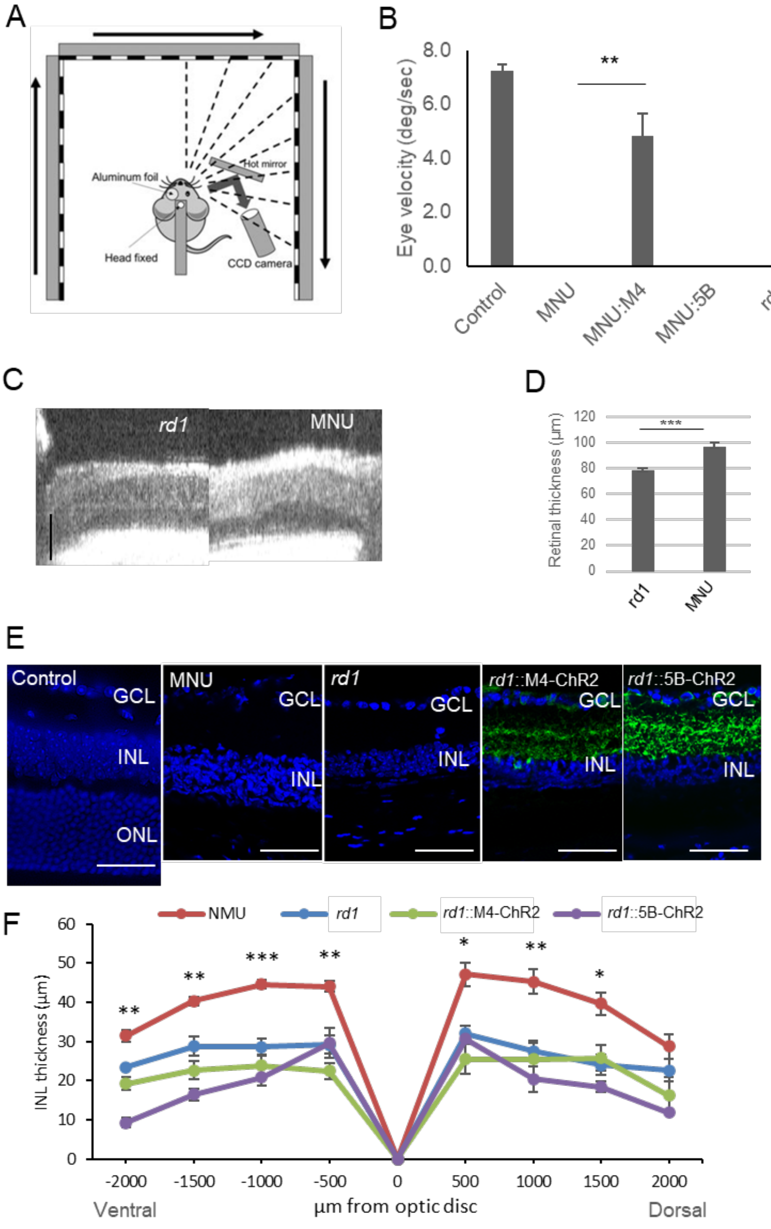


(A) Schematic view of the OKR system. The images of the right or left eyes are captured by a CCD camera placed on the same side. During measurement, the contralateral eyes are covered with aluminum foil. Visual stimulation is presented on three LCD monitors around the mouse, the head of which is fixed in the middle. (B) The average eye velocities of the control tetO-ChR2 mice (n = 3), MNU-injected tetO-ChR2 mice (n = 3), M4-ChR2M4-ChR2 mice (n = 9), 5B-ChR2 mice (n = 5)), *rd1*;tetO-ChR2 mice (n = 8), *rd1*;M4-ChR2M4-ChR2 mice (n = 10) and *rd1*;5B-ChR2 mice (n = 5) measured from the OKR system at 10 weeks of age. (C, D) Retinal structure images of *rd1* (n = 9) and MNU (n = 10)-treated mice at 8 weeks of age by optical coherence tomography (OCT). (E) Retinal sections of control (tetO-ChR2), MNU-treated tetO-ChR2, *rd1*::tetO-ChR2, *rd1*;M4-ChR2M4-ChR2 and *rd1*;5B-ChR2 mice at 8 weeks of age. Nuclear counter-staining with DAPI (blue) and ectopic gene induction with YC (green) are shown. (F) The INL thickness quantification from the sagittal sections. Cryosections from MNU-treated tetO-ChR2 (n = 6), and *rd1*::tetO-ChR2 mice (n = 6) at 8 weeks of age and *rd1*;5B-ChR2 (n = 3) and *rd1*;M4-ChR2M4-ChR2 (n = 3) mice at 10 weeks of age were used for quantification. (G) The YC-positive cells quantification in INL (cells per 200 x 200 μm section). *rd1*;5B-ChR2 (n = 3) and *rd1*;M4-ChR2M4-ChR2 (n = 3) mice at 10 weeks of age were used for quantification.

All scale bars: 50 μm. All error bars represent the SEMs. *$p < 0.05$, **$p < 0.01$, ***$p < 0.001$. Student’s 2-tailed t-test or Tukey's test.

**Figure S7. Gap junction was involved in the maintained response.**

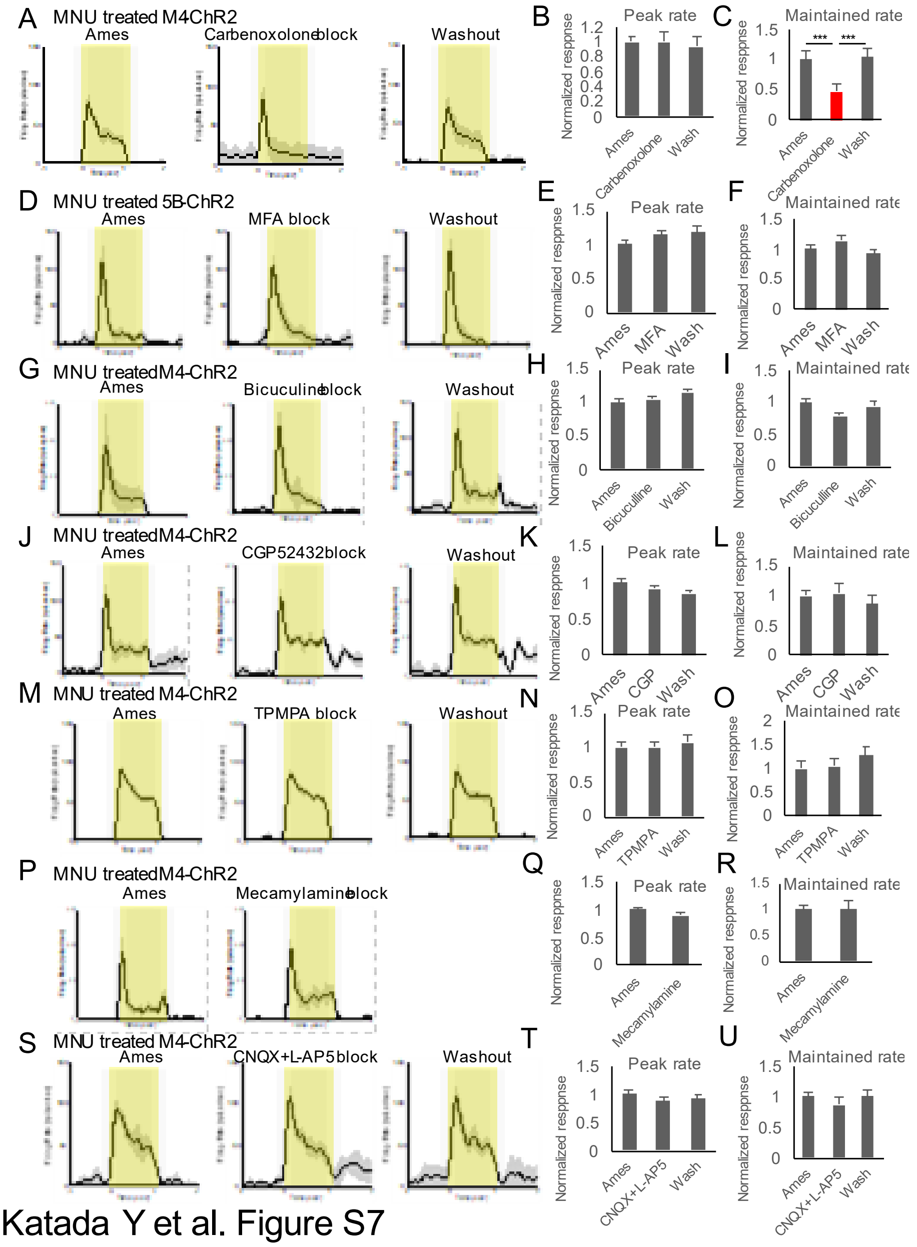


(A, D, G, J) Mean ± SEM of exemplar cell response firing rate recorded during normal Ames' medium superfusion (left), in synaptic block (middle), and after washout (right). MNU-treated M4-ChR2M4-ChR2 mice with carbenoxolone block (n=3 retinas, 25 cells)

(A), MNU-treated 5B-ChR2 mice with MFA block (n=3 retinas, 117 cells) (D), MNU treated M4-ChR2M4-ChR2 mice with bicuculline block (n=3 retinas, 148cells) (G), MNU treated M4-ChR2M4-ChR2 mice with CGP 52432 block (n=3 retinas, 130 cells) (J), MNU treated M4-ChR2M4-ChR2 mice with TPMPA block (n=3 retinas, 51 cells) (M), MNU treated M4-ChR2M4-ChR2 mice with mecamylamine block (n=3 retinas, 127 cells) (P), and MNU treated M4-ChR2M4-ChR2 mice with CNQX and L-AP5 block (n=3 retinas, 52 cells) (S). The gray areas around the averaged traces represent the SEM (B, C, E, F, H, I, K, L, N, O, Q, R, T, U). Averaged normalized peak firing rate and maintained rate. Maintained time frame is 0.4 to 1.0 seconds from light stimulation. Light intensity was 13.6 log photons/cm$^2$/s.

All error bars represent the SEMs. ***$p < 0.001$. One-way ANOVA and Tukey's test.

**Figure S8. Visual restoration profile of rAAV model**

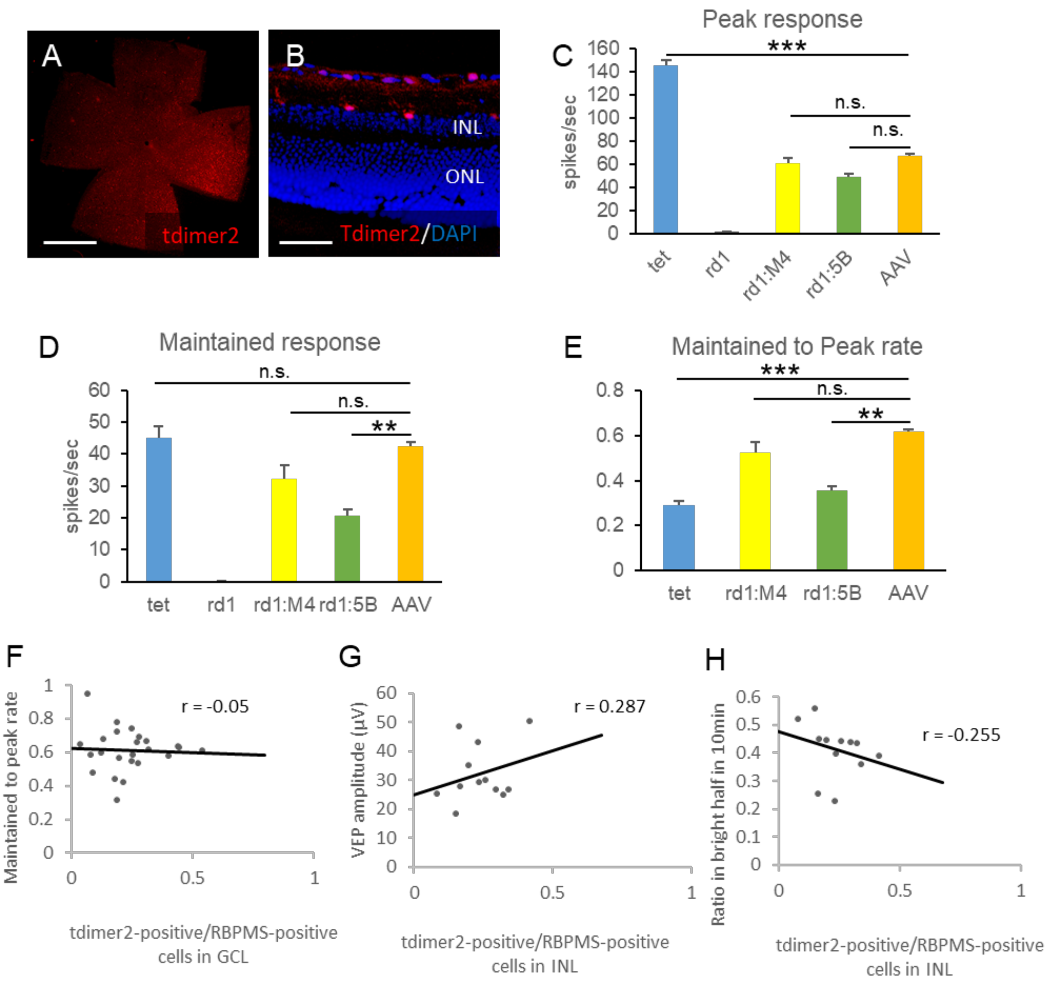


(A) AAV2–CAG–tdimer2–WPRE intravitreally injected mouse flat mounted retina and its section. (C-E) Comparison of peak response (C), maintained response (D) and maintained to peak rate (E) from MEA recordings among control (tetO-ChR2; n = 3 retinas, 112 cells), *rd1*::tetO-ChR2 (n = 3 retinas, 86 cells), *rd1*;M4-ChR2M4-ChR2 (n = 3 retinas, 18 cells), *rd1*;5B-ChR2 (n = 3 retinas, 17cells) and rAAV treated *rd1*::tetO-

ChR2 mice ($n = 24$ retinas, 1,151 cells) at 10 weeks of age. (F-H) Correlation between transfection efficiency into RGCs (tdimer2-positive cells/RBPMS-positive cells in INL) and maintained to peak rate (F) ($n = 24$), VEP amplitude (G) ($n = 24$) and % time in bright half in LDT (H) ($n = 12$). All error bars represent the SEMs. INL, inner nuclear layer; GCL, ganglion cell layer, RGC, retinal ganglion cell, ONL, outer nuclear layer. Scale bars, 1,000 μm in (A), 50 μm in (B), n.s.: not significant, *$p < 0.05$, **$p < 0.01$, ***$p < 0.001$. Games-Howell test (C, D, E), Pearson's correlation coefficient (F-H).
